# A Most Powerful Test for Gene–Gene Interaction in the Presence of Main Effects

**DOI:** 10.64898/2026.01.30.702572

**Authors:** Razvan G Romanescu, Michelle Liu

**Affiliations:** University of Manitoba

## Abstract

We consider the problem of optimal testing for genetic interaction between two variants, allowing for possible main effects. Finding a most powerful test is important because it ends a series of attempts in the literature to construct ever more powerful tests for interaction at the variant pair level. Testing under a logistic regression model is known to be underpowered, partly because patterns of enrichment in the genotypes themselves are lost when regarding genotypes solely as predictors. Instead, we use the retrospective likelihood approach, which makes use of all the data by treating genotypes as outcomes alongside affection status. Using a parsimonious parameterization of penetrance based on the risk ratio, which links directly to the population prevalence and avoids having to estimate an intercept term, we construct an approximate uniformly most powerful unbiased test for interaction. This test is based on optimal testing theory and accounts for nuisance main effects without requiring their explicit estimation. The test statistic can be easily modified for optimal testing under other modes of genetic interaction, such as recessive × recessive or dominant × dominant. We demonstrate significant power gains compared to the odds–ratio–based PLINK test, in simulation studies. Finally, we apply the test to scan for interactions in IBD cases and controls from the UK Biobank. The top SNP pairs show enrichment for a pathway related to existing therapies for IBD.

## 1 Introduction

The genome-wide association study (GWAS) era has produced a fairly complete description of main effects for common genetic variants. Even relatively small effect sizes have been mapped for most complex diseases in large panel studies of unrelated individuals. However, marginal effects explain a relatively low proportion of estimated heritability. One important source of “missing heritability” is thought to be higher order genetic effects, in particular genetic interaction, broadly termed as epistasis (Jung et al., 2023; Stringer et al., 2013). In a statistical inferential sense, epistasis refers to the joint effect of two genetic variants that contributes to risk beyond the cumulation of their individual marginal effects. Computationally, the probelm of testing for genetic interaction has been difficult due to the large number of tests involved, and therefore, the stringent multiple testing corrections applied.

The most widely–used parametric risk model that includes interaction effects is logistic regression with disease state as outcome, genetic dosages of variants *A* and *B* as main effects, and the dosage product as interaction. This is a standard risk model in biostatistics and has the advantage that it can include covariates as additional linear terms. However, this is not the most powerful model, partly because it has a number of parameters that need to be estimated, but more importantly because it does not explain the frequency pattern in genotype combinations, which is simply assumed as a fixed input. This pattern contains valuable information for detecting interactions, and there are non–parametric case–only tests that leverage this information to detect interaction signals. By contrast, one would not be able to fit a logistic regression in the absence of controls. The method that models genetic dosages specifically is the retrospecive likelihood approach (Bhattacharjee et al., 2010; Chatterjee & Carroll, 2005). By using genetic dosages themselves as outcomes, in addition to the affection status, thereby increasing the effective sample size. This paper starts from the restrospective likelihood for the case-control design, and seeks to find the most powerful test for interaction, in the presence or absence of main effects. Unlike the logistic regression formulation, where interaction is simply added on as an extra linear term to pre-existing main effects, we treat the main effects as nuisance parameters and do not seek to estimate them, but only adjust for them to keep type I error control nominal.

The theory of uniformly most powerful tests in the single parameter case is well–known, and part of traditional statistical education. Extending optimality results to the multi– parameter model, where one parameter is of interest and the others are nuisance, is more conceptually involved. Even though it is also part of classical statistical inference theory (Lehmann & Romano, 2008) its uptake in practice has been limited at best, possibly because of its relatively high level of abstraction and because it often requires the extra assumption of test unbiasedness. However, there are some strong reasons why optimal testing theory can benefit methods in statistical genetics, in particular. First, as power is top of mind in genome–wide scans, finding a most powerful test not only ensures it is more powerful than competing tests, but also establishes the power envelope as a benchmark against which to compare other tests. This closes the door to further research on more powerful tests for the postulated risk models, enabling methodologists to focus on better modeling of disease risk – such as researching mechanisms of action at the pathway level – as this will more likely close the gap between statistical testing and a biological understanding of disease aetiology. Secondly, there are many situation in genetics when the risk model contains parameters the investigator is not interested in, such as when adjusting for covariates. Typically, the options are either to assume their effect is negligible, and hence to ignore them, or to estimate them. Optimal testing theory in the presence of nuisance parameters gives a third option, namely to condition them out of inference. The constructed test statistic is guaranteed to be uniformly most powerful unbiased (UMPU) whatever the values of the nuisance parameters. This is certainly a desirable property to have if one wishes to disentangle interactions from other effects.

## 2 Methods

### 2.1 Basic setup and notation

The data can be tabulated in a 3 × 3 table of genotypes corresponding to dosages of 0, 1, and 2 risk alleles for each of two variants, A and B. Denote by *x*_*ij*_, *i, j* ∈ {0, 1, 2}, the number of cases having *i* alleles A and *j* alleles B (out of a total of *n*_1_ ascertained cases). Similarly, *y*_*ij*_ counts the controls. The distribution of vectors **x** and **y** is multinomial, with totals *n*_1_ and *n*_0_. To obtain the genotype probabilities *P* (*i, j* | case) and *P* (*i, j* | control) for each genetic combination (*i, j*), we use Bayes’ law:

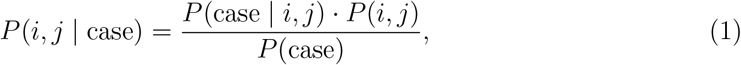

where *P* (*i, j*) is the probability of observing genotype (*i, j*) in the general population, which can be computed from the minor allele frequencies of A and B, *µ*_*A*_ and *µ*_*B*_. For example, if we denote *P* (*i, j*) = *µ*_*ij*_ then, assuming Hardy-Weinberg equilibrium we find *µ*_00_ = (1 − *µ*_*A*_)^2^(1− *µ*_*B*_)^2^, etc. Next, *P* (case) is the unconditional probability of being affected which, by definition, is the prevalence of condition *D* in the population and is given by parameter *π*. The last term, *P* (case *i*, | *j*), is the penetrance of genotype (*i, j*). In what follows, let us assume that the penetrance can be described as a function of one unknown parameter, *γ*, which controls the effect size *P* (*D* = 1 | *i, j*) = *f* (*γ*; *i, j*). In the next section, we will further specify function *f*. For now, we can rewrite formula (1) as

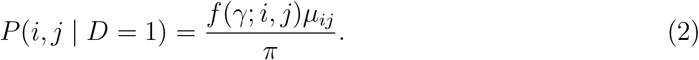

Similarly for controls:

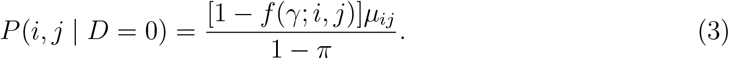

Using (3) and (4), the retrospective likelihood function of all the data, *P* (***x, y***| *γ*), is computed using multinomial probabilities as

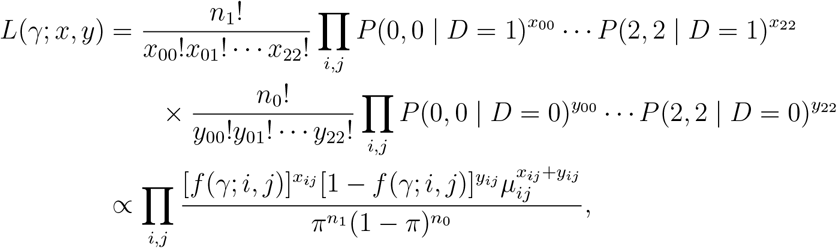

with log-likelihood:

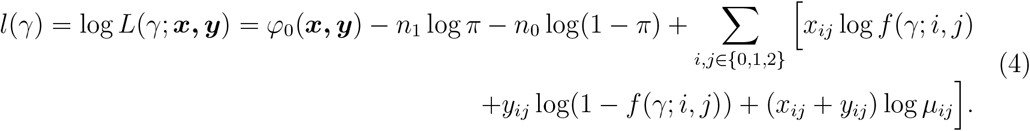

Here, we regard *γ* as the unknown parameter and assume the other parameters, *π, µ*_*A*_, and *µ*_*B*_, to be known. This is reasonable since they are population level quantities usually available from the literature (*π*), or that can be queried from large panels of genetic variation (*µ*_*A*_ and *µ*_*B*_).

### 2.2 The Multiplicative Interaction Model

The model of disease risk we use is multiplicative in genetic main effects and interaction for two loci, in addition to other covariates. The form of the model is loosely based on Hu et al. (2014) for the genetic part. We define the penetrance function as:

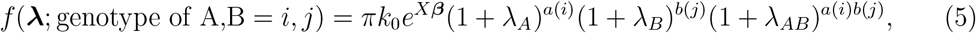

where *k*_0_ is a normalizing constant, *X* is a matrix of covariates, and the two variants *A* and *B* have effect sizes *λ*_*A*_, *λ*_*B*_ and interaction effect *λ*_*AB*_. A default choice for functions *a* and *b* is the identity function;this makes the exponent of the penetrance similar to the linear log–odds formulation in logistic regression. We will call this model “additive” in both variants, following the nomenclature in Ueki & Cordell (2012), which will be assumed for most of the paper. However, it is easy to accommodate dominant and recessive effects in either variant by turning *a* and *b* into step functions. Note that constant *k*_0_ = *k*_0_(***λ, µ***, *X*) is computed to make the overall penetrance for the population equal to *π* (see derivation in the Supplementary Material). This formulation is intuitive because the risk of disease for an individual who has all covariates equal to zero is *πk*_0_, and every additional non-zero covariate contributes multiplicatively to risk, starting from this baseline. This model is an extension of the Risch risk model, introduced in (Risch, 1990; Risch & Merikangas, 1996) and used in (Lu & Elston, 2008; Pharoah et al., 2002; Slatkin, 2008; Wray & Goddard, 2010; Wray et al., 2007) as a risk model for multilocus diseases. The difference with our formulation (5) is that, while in the Risch model all mutations across multiple loci are added together and have the same effect, we allow for distinct effects for both the two main effects as well as their interaction.

This model has some notable differences to the more traditional and widely–used logistic regression risk model with main effects and interaction (see Ueki and Cordell, 2012, and others). While in the latter, the odds ratio is easy to obtain in terms of the parameters, in this model it is more natural to obtain relative risk, or risk ratio for one extra allele *A*, as 1 + *λ*_*A*_. Similarly for the interaction, i.e. the risk of one extra allele for both *A* and *B* over the marginal effect for each, is 1 + *λ*_*AB*_. Formulation (5) is also preferred because it provides a more mathematically transparent way to link individual risk to population prevalence *π*.Using an external estimate for *π* saves estimating one parameter, whereas in logistic regression it is not easy to calibrate to *π* in closed form, and, as a result, we are stuck with having to estimate the intercept, which costs power and is not of interest.

Before proceeding, it is worth asking the question of how our definition of interaction is related to the concept of epistasis, which is what we are ultimately hoping to measure. According to Phillips (2008), the concept of epistasis, or interaction among genes can be viewed in three ways: (i) functional epistasis – referring to the molecular level interaction; (ii) compositional epistasis, i.e., the blocking effect of an allele at one locus on an allele at a different locus and (iii) statistical epistasis – the statistical interaction which adds a non-linear effect to the additive genetic model. Statistical interaction, visible in the penetrance model above via the product of coded dosages *a*(*i*)*b*(*j*), should not be viewed as yet another definition of interaction, but rather exists to support (i) and (ii), making it a natural modeling choice for most generic tests for interaction. In particular, this model is apt at describing (i) by drawing a parallel to the speed of a chemical reaction where reagents can be thought of as the protein products of the two genes, with the genetic dosages of the loci in question being the “concentrations” of the substances reacting. In this case, the concentrations have a multiplicative effect on the speed, or quantity of the product (i.e., the interaction effect) at a specific time during the reaction, as is typical in chemical kinetics. The effect of interaction in (ii) is, perhaps, better modeled by an on/off threshold for one of the variants’ dosages, which can be specified via a dominant/recessive coding for one of the variants (the blocking one).

## 3 Optimal statistical tests

In this section, we look to construct optimal tests for interaction for the risk model defined previously. Ultimately, the aim is to develop a uniformly most powerful unbiased test for epistatic effects, which is well controlled for type I error whether there are main effects or not.

### 3.1 Locally most powerful test in the absence of main effects

With no main effects or other covariates, penetrance (5) becomes:

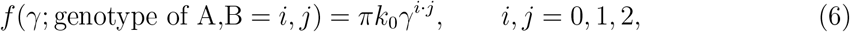

where *γ* is the size of the interaction effect (*γ* = 1 + *λ*_*AB*_), the unique argument of the penetrance function in (2 –3). Under the null of no association, *k*_0_ = 1, the penetrance *f* ≡ *π*, and expected genotype fractions are the same as in the general population, namely *P* (*i, j* | *D* = 0 or 1) = *µ*_*ij*_. Under the alternative *H*_*a*_: *γ* <> 1 we have *k*_0_ >< 1 and the expected number of cases and controls for each genotype combination is multiplied by a factor given in Table 3.1, relative to the general population.

**Table 1:** Relative abundance or scarcity of genotype combinations in cases and controls.

| $P(\text{Genotype} \mid D = 1)/P(\text{Genotype})$ | A=0 | 1 | 2 |
| --- | --- | --- | --- |
| B=0 | $k_0$ | $k_0$ | $k_0$ |
| 1 | $k_0$ | $k_0\gamma$ | $k_0\gamma^2$ |
| 2 | $k_0$ | $k_0\gamma^2$ | $k_0\gamma^4$ |

| $P(\text{Genotype} \mid D = 0)/P(\text{Genotype})$ | A=0 | 1 | 2 |
| --- | --- | --- | --- |
| B=0 | $\frac{1-\pi k_0}{1-\pi}$ | $\frac{1-\pi k_0}{1-\pi}$ | $\frac{1-\pi k_0}{1-\pi}$ |
| 1 | $\frac{1-\pi k_0}{1-\pi}$ | $\frac{1-\pi k_0\gamma}{1-\pi}$ | $\frac{1-\pi k_0\gamma^2}{1-\pi}$ |
| 2 | $\frac{1-\pi k_0}{1-\pi}$ | $\frac{1-\pi k_0\gamma^2}{1-\pi}$ | $\frac{1-\pi k_0\gamma^4}{1-\pi}$ |

When testing two–sided hypotheses, the likelihood ratio test (LRT) can be shown to be approximately UMP unbiased. The test statistic is 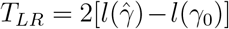, which is distributed as *χ*^2^ with 1 degree of freedom. With this assumed penetrance function, the LRT becomes (see the Supplementary Material for details)

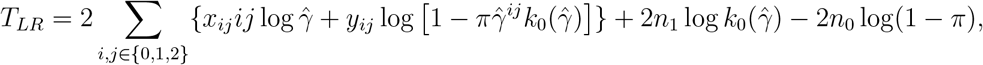

where 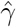 is the unrestricted MLE of *γ*. As we shall see in simulation studies, the power profile of this test is about as good as one can hope for, however its type I error control can be off when main effects are present.

## 4 Derivation allowing for main effects

The additive version of penetrance model (5) with main effects, interaction and no covariates reads

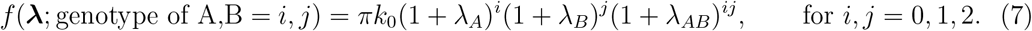

One might be tempted to think that extending the LRT to the multivariate case of three parameters would provide the desired solution, where the likelihood is optimized over two parameters (*λ*_*A*_ and *λ*_*B*_) under the null and over all three under the alternative hypothesis. However, optimal testing theory does not, in fact, extend to the multivariate case as easily. Although the LRT described would be a legitimate test, there is no guarantee that it is UMPU over all possible values of *λ*_*A*_ and *λ*_*B*_; for some combinations of true effect sizes, another test may be more powerful. Theory does exist, though, for constructing a UMPU test for a parameter of interest in the presence of other, nuisance parameters. For a test to be most powerful in this setting, we need to show that it is most powerful given any value of the sufficient statistics for the nuisance parameters.

We start by obtaining the sufficient statistics for model (7). The log likelihood for this genetic model can be obtained by developing (4) and using an approximation for small effect sizes. We do this to keep the mathematics tractable; consequently, the conclusions of this development will be for a locally most powerful test (due to the approximations around zero effects). Thus, for unknown parameter vector ***λ*** = (*λ*_*A*_, *λ*_*B*_, *λ*_*AB*_), we have

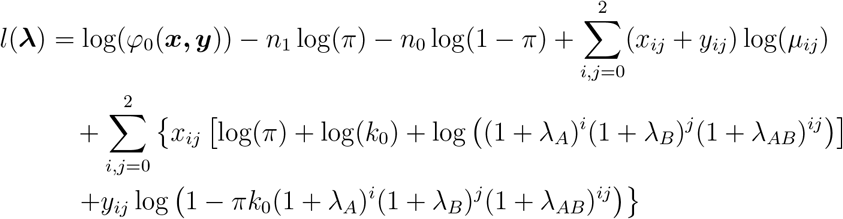

Making use of the fact that effect sizes ***λ*** are small, we can approximate (1 + *λ*_*A*_)^*i*^(1 + *λ*_*B*_)^*j*^(1 + *λ*_*AB*_)^*ij*^ *≈* 1 + *iλ*_*A*_ + *jλ*_*B*_ + *ijλ*_*AB*_. Similarly, terms of the form log(1 − *π*(1 + *δ*)) can be approximated by 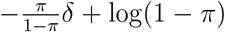, for *δ ≈* 0. For instance, log(1 − *πk*_0_) can be approximated in two steps, using the value of *k*_0_ from (2) in the Supplementary Material:

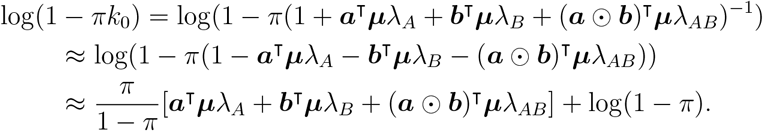

Here, constant vectors ***a*** and ***b*** are defined as

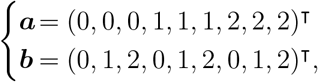

where ⊙ is the Hadamard (or element-wise) vector product. As an aside, weight vectors ***a*** and ***b*** not only make the math more compact, but also encode the genetic model, such that switching to a different model only requires changing ***a*** and ***b***. The order of weights correspond to the ordering of genotypes in data vector ***x*** = (*x*_00_, *x*_01_, *x*_02_, *x*_10_, *x*_11_, *x*_12_, *x*_20_, *x*_21_, *x*_22_)^⊺^, which is also followed in ***y*** and ***µ***. With these approximations, the log likelihood function can be written as

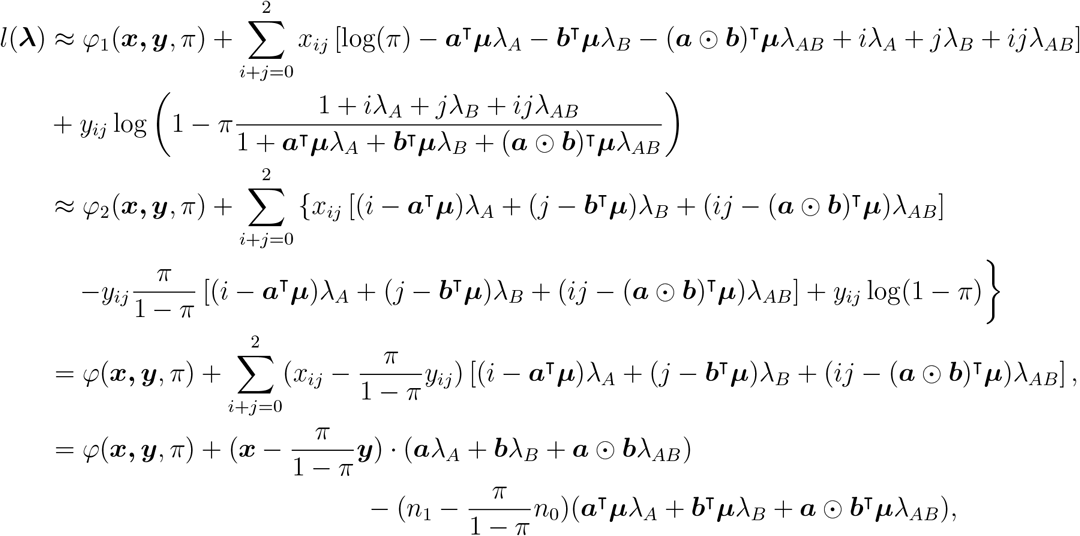

Putting 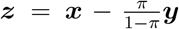, we can rewrite the log likelihood in the canonical form of the exponential family for a distribution with parameters *λ*_*A*_, *λ*_*B*_, and *λ*_*AB*_:

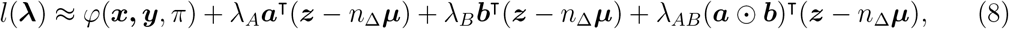

where we have denoted 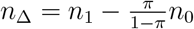. Defining the new summary statistics

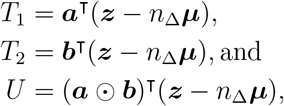

we can deduce from (8) that (*U, T*_1_, *T*_2_) are complete, sufficient statistics for parameter vector (*λ*_*A*_, *λ*_*B*_, *λ*_*AB*_) in (***X, Y***).

Our interest is in testing for *λ*_*AB*_, so we treat *λ*_*A*_ and *λ*_*B*_ as nuisance. From (8), we can see that on the boundary between the null and alternative hypotheses, i.e., when (*λ*_*A*_, *λ*_*B*_, *λ*_*AB*_) ∈ ℝ^2^ ×{0}, (*T*_1_, *T*_2_) are the sufficient statistics of a two parameter exponential family, and thus contain all the information available for inference on *λ*_*A*_ and *λ*_*B*_. By definition, then, the conditional distribution of ***z***|*T*_1_, *T*_2_ should be independent of *λ*_*A*_ and *λ*_*B*_, thus, parameter-free on the boundary. Similarly for that of *U* |*T*_1_, *T*_2_. The strategy here is that after conditioning on the effect of the nuisance variables, the resulting test for *λ*_*AB*_ will be optimal regardless of what the true *λ*_*A*_ and *λ*_*B*_ are. This will be made more clear in subsection 4.2. Before we can proceed, we require a distribution of the test statistics, which all depend on ***z***. We model this variable next.

### 4.1 Distributional characterization of the statistic z

We wish to calculate the distribution of ***z*** under the null (i.e., when *λ*_*AB*_ = 0). We know that ***x*** = (*x*_00_, *x*_01_, …, *x*_22_) is distributed as Multinomial(***p***, *n*_1_), where

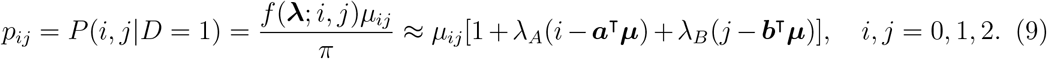

As such, the mean of *X*_*ij*_ is *E*[*X*_*ij*_] = *n*_1_*p*_*ij*_, the variance Var[*X*_*ij*_] = *n*_1_*p*_*ij*_(1 − *p*_*ij*_) and the pairwise covariances are 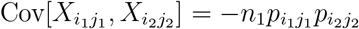. These can all be written in terms of *λ*_*A*_ and *λ*_*B*_, using the expression for *p*_*ij*_ from above (see derivation in the Supplementary Material). We can then approximate the distribution of ***x*** via a multivariate normal, which matches the mean vector and covariance matrix of the multinomial, i.e.,

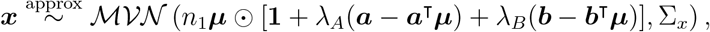

where we use the notation ***a*** − *c* with *c* scalar to mean ***a*** − *c***1**. Similarly, (*y*_00_, *y*_01_, …, *y*_22_) *∼* Multinomial(***q***, *n*_0_), where

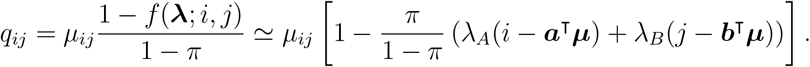

We make the same approximation via the multivariate normal:

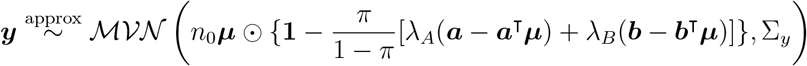

We can now obtain the distribution of vector 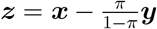. Its mean is easy to obtain; the variance-covariance matrix is given by 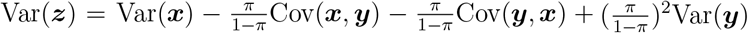. Using the fact that ***x*** and ***y*** are independent samples, the covariance terms are zero matrices, and we have

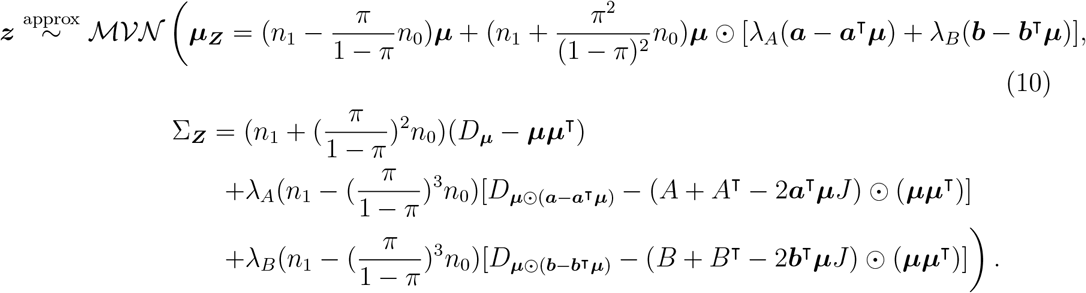

We can also write the mean and variance of ***z*** more compactly as

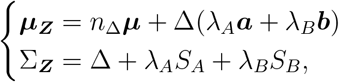

where matrix 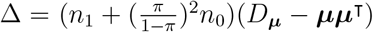, and matrices *S*_*A*_ and *S*_*B*_ can also easily be identified from (10).

### 4.2 A most powerful test for *λ*_*AB*_ in the presence of nuisance main effects

As mentioned before, inference theory provides a way to build a most powerful test in the presence of nuisance parameters by using *U*| ***T***, i.e., the conditional distribution of sufficient statistics. In practical applications, however, such conditional distributions may be cumbersome to obtain in closed form; for this reason, a simplified version of this approach has been proposed in (Bhattacharya & Burman, 2016) and (Lehmann & Romano, 2008). This is based on finding a pivotal quantity that is a function of only *U* and ***T*** and has a parameter-free distribution under the null. Thus, let *V* be defined as

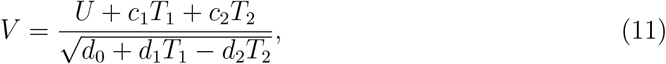

where *c*_1_, *c*_2_, *d*_0_, *d*_1_, *d*_2_ are constants to be specified. We first show that V is a pivotal quantity.

#### Lemma 1.

The distribution of *V* is approximately standard normal for appropriately chosen constants *c*_1_, *c*_2_, *d*_0_, *d*_1_, and *d*_2_.

PROOF: Starting from the definitions of *T*_1_, *T*_2_ and *U*, the numerator can be written as (***a*** ⊙ ***b*** + *c*_1_***a*** + *c*_2_***b***)^⊺^(***z*** − *n*_Δ_***µ***), which is distributed as

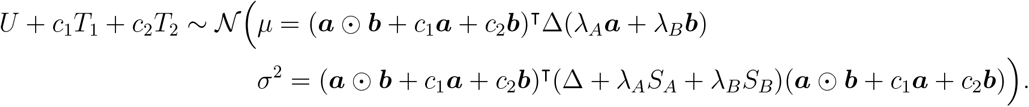

We can choose *c*_1_ and *c*_2_ to make the mean identically zero, namely

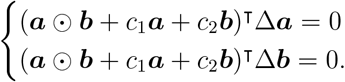

The variance of the numerator becomes

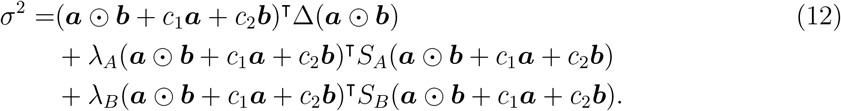

We can choose the coefficients *d*_0,1,2_ to make the sum in the denominator on average *σ*^2^, namely

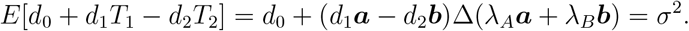

Matching the coefficients of *λ*_*A*_, *λ*_*B*_ and the constant term gives us

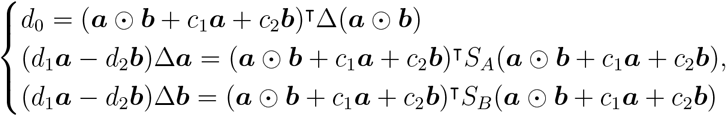

or

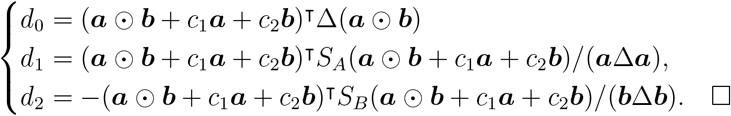

Next, we show that a test based on *V* is most powerful and unbiased for *λ*_*AB*_, in the presence of nuisance parameters. We remind the reader that an “unbiased test” is one that has power to detect less than or equal to *α* at all points in the null parameter space, and at least *α* at all points in the alternative space. This condition avoids tests that are only powered for one–sided effects and lack any power for effects with the opposite sign.

Theorem 1. The test *φ*(*V*) = *I*{*V* < *z*_*α/*2_ or *V* > *z*_1−*α/*2_}, for *V* as in Lemma 1, is approximately UMPU for testing *H*_0_: *λ*_*AB*_ = 0 vs. *H*_1_: *λ*_*AB*_ ≠ 0.

PROOF: For the problem of two-sided testing, (Bhattacharya & Burman, 2016) require the following conditions for a test *φ* based on a quantity *V* = _*g*_(*U*, ***T***) to be UMP unbiased at level *α*:

a. *V* is independent of ***T*** when *λ*_*AB*_ = 0, and
b. *g* can be written *g*(*u*, ***t***) = *a*(***t***)*u* + *b*(***t***), where *a*(***t***) > 0.

Furthermore, the rejection region of test *φ*(*v*) = *I* {*V* < *c*_*l*_ or *V* > *c*_*u*_} has to satisfy (for continuous random variables) the following conditions:

i. 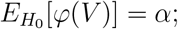
ii. 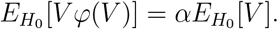

In particular, condition (i) controls nominal type I error, and (ii) is implied by unbiasedness. It is easy to check that, for *V* being standard normal, both of the latter conditions hold for symmetric rejection regions. Condition (b) is also trivial to see.

To prove (a), we will make use of Basu’s theorem (Lehmann & Romano, 2008), which says that if the family of distributions of ***T*** is boundedly complete, and if *V* is any ancillary statistic – by which we mean any statistic whose distribution does not depend on the parameters of the model –, then *V* is independent of ***T***. This is the case under the null, when our probability model can be described by parameter vector (*λ*_*A*_, *λ*_*B*_), for which ***T*** is sufficient. ***T*** = (*T*_1_, *T*_2_) is also complete, which loosely means that its dimension matches the dimension of the parameter space, in this case two. As completeness implies bounded completeness, the first part of Basu’s theorem is satistied. By Lemma 1, *V* is an ancillary statistic, which completes the proof. This reasoning is spelled out more explicitly in Corollary 5.1.1 in (Lehmann & Romano, 2008) for the case of an exponential family in which one parameter is fixed (in our case, *λ*_*AB*_ = 0), the conclusion being the same, that *V* is independent of ***T***.

Thus, our test *φ* is approximately UMPU for testing *H*_0_: *λ*_*AB*_ = 0 vs. *H*_1_: *λ*_*AB*_ ≠ 0, with the approximation stemming largely from the fact that (4) is a linearization for small effect sizes. □

### 4.3 Testing for other models of interaction

Having done all the work above for the additive model, extending the test to other models of genetic interaction is straightforward. The only change needed for using the conditional test is updating vectors ***a*** and ***b*** to encode the new penetrance for all nine genotype combinations. Table 2 shows the penetrance formulations and coding for the models recessive × recessive, dominant × dominant, and dominant × additive. These have been described in (Ueki & Cordell, 2012). In all these cases, coding for the interaction effect is given by ***a*** ⊙ ***b***. All results so far hold in the same way using the updated coding.

**Table 2:** Alternative genetic interaction models, including how they are coded in the corresponding UMPU test statistic. *k*_0_ is chosen to make the overall population penetrance equal to *π* under each model.

| Model of interaction | Penetrance function<br>$f(\boldsymbol{\lambda}; \text{genotype of A,B} = i, j) = \pi k_0 \times \dots$ | $\mathbf{a}$ and $\mathbf{b}$ coding |
| --- | --- | --- |
| Recessive x Recessive | $(1 + \lambda_A)^{I\{i=2\}}(1 + \lambda_B)^{I\{j=2\}}(1 + \lambda_{AB})^{I\{i=j=2\}}$ | $\mathbf{a} = (0,0,0,0,0,0,1,1,1)$<br>$\mathbf{b} = (0,0,1,0,0,1,0,0,1)$ |
| Dominant x Dominant | $(1 + \lambda_A)^{I\{i>0\}}(1 + \lambda_B)^{I\{j>0\}}(1 + \lambda_{AB})^{I\{ij>0\}}$ | $\mathbf{a} = (0,0,0,1,1,1,1,1,1)$<br>$\mathbf{b} = (0,1,1,0,1,1,0,1,1)$ |
| Dominant x Additive | $(1 + \lambda_A)^{I\{i>0\}}(1 + \lambda_B)^j(1 + \lambda_{AB})^{I\{i>0\}j}$ | $\mathbf{a} = (0,0,0,1,1,1,1,1,1)$<br>$\mathbf{b} = (0,1,2,0,1,2,0,1,2)$ |

**Table 3:**
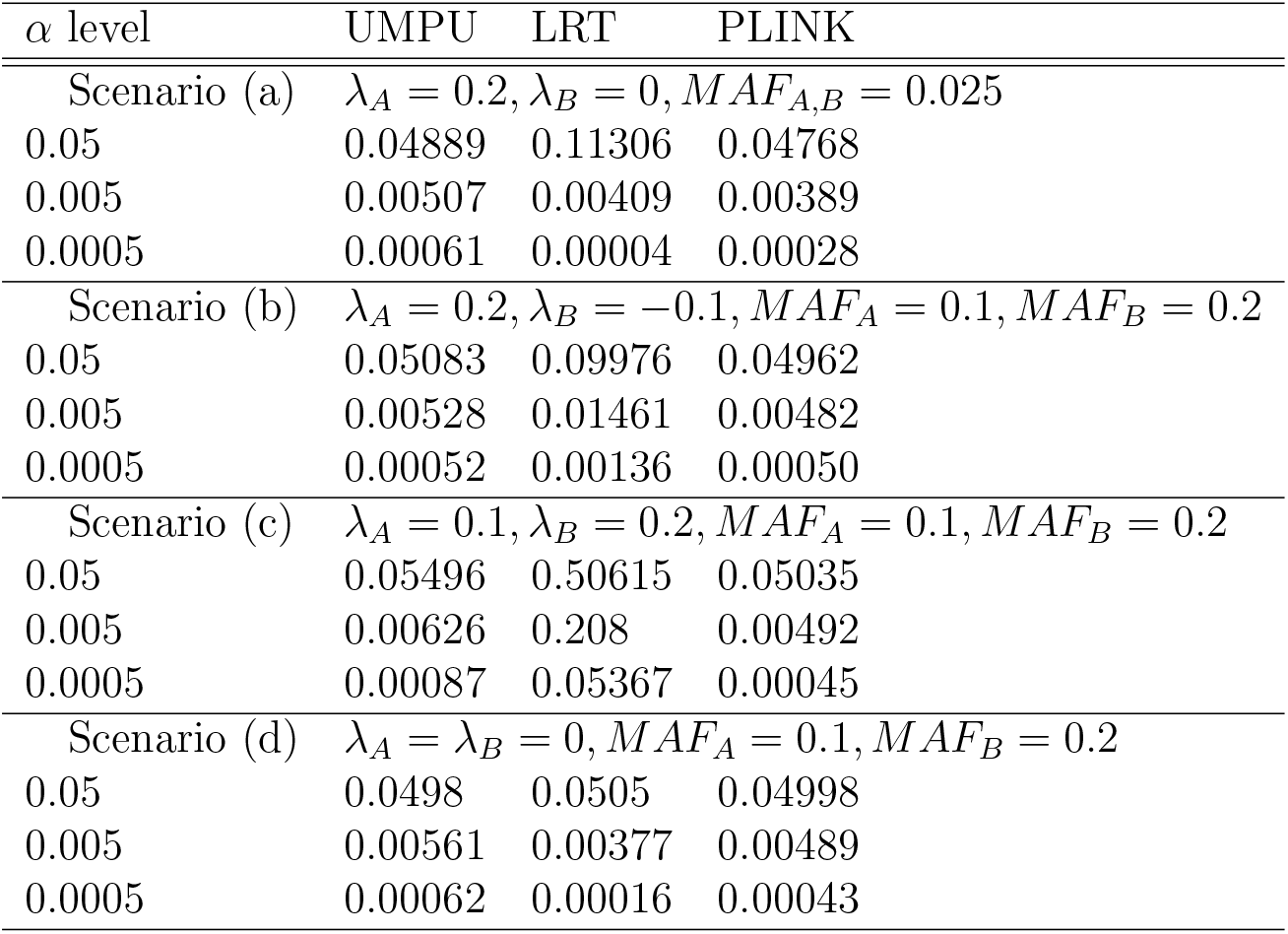
Type I error rates in four different simulation scenarios for the additive × additive model. Uses study sizes *n*_1_ = *n*_2_ = 10, 000 for scenario (a) and *n*_1_ = *n*_2_ = 1, 000 for the other scenarios.

### 4.4 Other tests

We will compare our results against the original PLINK *fast-epistasis* test. This test has been described in detail elsewhere (Ueki & Cordell, 2012). Briefly, this test tabulates genotypes for the two variants A and B into a 2 by 2 table, by counting the total number of reference and alternative alleles combinations a/A and b/B in each cell. Calling the four counts in the table of cases *a, b, c*, and *d*, it estimates the odds ratio *R* = *ab/cd* with variance Var[log(*R*)] = 1*/a* + 1*/b* + 1*/c* + 1*/d*. Similarly for controls, it estimates a different odds ratio *S*. Finally, a standardized difference of the log odds of cases versus the log odds of controls is computed as

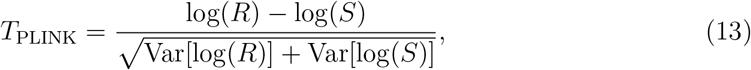

which follows a standard normal distribution. Besides using four genotype combinations compared to our nine, another difference is that the PLINK test does not fully rely on the independence assumption of variants under the null. In other words, it will still work when loci A and B are correlated in unaffected individuals, such as when the variants are in LD, because the test looks for a difference in pattern between cases and controls. By contrast, the case–only version of the PLINK test, as well as our test, would detect variants in strong LD as significant, due to their departure from independence. For completeness, we should mention that the current implementation in PLINK of the epistasis test might be slightly different from the one described above. When we refer to the ‘PLINK’ test we mean specifically the test defined in (13), which also follows the description in the PLINK online manual v.1.07 for the `--fast-epistasis` command.

Although there are a number of other tests at the variant level, including Wu et al. (2010) and others as reviewed in Ueki & Cordell (2012) and Evangelou (2018), we do not compare against them for two reasons. Firstly, many of these tests have been compared before, including with the PLINK test. Ueki & Cordell (2012) report that, in the case of the additive × additive model (the most common case), only the original Wu test stands out from the rest, however, when adjusted for type I error, its performance is relatively average. The PLINK test also has an average (thus competitive) power profile, making it a good benchmark, given that it is also the simplest test. Secondly, we have proved our test to be UMPU, thus, for our assumed penetrance models we no longer need to compare power with existing tests; comparison against PLINK is only done to give an idea of expected power gains.

Testing in the most recent literature focuses on two directions with significant overlap: (i) machine learning (ML) based methods such as BOOST (Wan et al., 2010), GenEpi (Chang et al., 2020), and multifactor dimensionality reduction (Hou et al., 2019; Park et al., 2021; Xu et al., 2016); and (ii) multi–locus or group testing methods (including GenEpi; (Emily, 2016; Guo et al., 2021a,b; Xia et al., 2015) and others). ML methods are often built for speed of computation and may not require an explicit risk model for each variant. The problem in general with ML tests is that, due to their generality, they cannot typically guarantee the most power for a specific risk model (e.g., the BOOST test is more powerful than PLINK in some situations, but less so in others (Wan et al., 2010)). Thus, showing that a test is UMPU makes it state–of–the–art, as long as no other competing methods have the same property. Set or pathway tests for interaction are particularly sensible due to reducing the overall number of tests, and this area is likely to become the future of interaction testing. Even here, our blueprint for obtaining a parametric UMPU test may be extended for a multi–locus penetrance and the resulting test is likely to outperform existing set–based interaction tests, at least for the assumed risk model.

## 5 Simulation study

### 5.1 Setup

In this section we generate two unlinked genetic variants for a large population pool of 10, 000, 000 unrelated individuals. For each individual, we independently generated the gene dosage for the two SNPs (A and B), according to minor allele frequencies MAF_*A*_ and MAF_*B*_. Based on their genotype, we simulate each individual’s disease risk according to model (7). The baseline population prevalence for the disease was set at 10% (*π* = 0.1), and the main and interaction effects (*λ*_*A*_, *λ*_*B*_, and *λ*_*AB*_) are varied across simulation scenarios.

For one study we ascertain *n*_1_ cases and *n*_0_ controls from the population pool, according to their disease risk, i.e., we sample individuals sequentially until both groups are filled. Each test is computed using the genotype data of the cases and controls, assuming the known population minor allele frequencies. Type I error and power are computed empirically for *λ*_*AB*_ = 0 versus *λ*_*AB*_ ≠ 0, using 100,000 replications. We consider significance levels × 10^−2^, ×10^−3^ and ×10^−4^ for type I error, and *α* = 5 × 10^−4^ for power.

### 5.2 Results

Table 3 shows the empirical type I error estimates in four scenarios, including with and without main effects, common and uncommon variants. The study sizes used were *n*_1_ = 1, 000 cases and *n*_0_ = 1, 000 controls in all cases except the low–frequency case (scenario a), where *n*_1_ = *n*_0_ = 10, 000. We can see that the conditional (UMPU) and PLINK tests attain decent type I error control overall, whereas the LRT’s performance depends on the situation. In certain cases type I errors for the LRT can be much higher or lower than nominal.

Note that the LRT and UMPU are as developed under an additive × additive interaction model, as this is likely what an analyst would use as a general-purpose scanning tool. We then look at data simulated under each of the models in Table 2. Again, studies of sizes *n*_1_ = 1, 000 cases and *n*_0_ = 1, 000 controls are ascertained, and we analyze them using the same three tests. We also include a specialized conditional UMPU test for the correctly specified genetic model, in order to show its performance. Table 4 shows that, generally, type I error is well controlled for both versions of the UMPU, as well as PLINK, whereas departures from nominal can be significant with the LRT.

**Table 4:** Type I error rates in four different simulation scenarios for the dominant and recessive models. The UMPU and LRT tests are misspecified (i.e., do not reflect the simulation model), while the ‘UMPU–specialized’ test is correctly specified. Uses study sizes of *n*_1_ = *n*_2_ = 1, 000.

| $\alpha$ level | UMPU | UMPU–<br>specific | LRT | PLINK |
| --- | --- | --- | --- | --- |
| Scenario (e) | $D \times D, \lambda_A = 0, \lambda_B = -0.1, MAF_A = 0.1, MAF_B = 0.2$ | | | |
| 0.05 | 0.05164 | 0.05041 | 0.08527 | 0.05077 |
| 0.005 | 0.00582 | 0.00559 | 0.00754 | 0.00494 |
| 0.0005 | 0.00066 | 0.00058 | 0.00019 | 0.00052 |
| Scenario (f) | $D \times A, \lambda_A = 0.2, \lambda_B = -0.1, MAF_A = 0.1, MAF_B = 0.2$ | | | |
| 0.05 | 0.04956 | 0.0511 | 0.07723 | 0.04876 |
| 0.005 | 0.00484 | 0.00526 | 0.00943 | 0.00467 |
| 0.0005 | 0.00047 | 0.00048 | 0.00079 | 0.0004 |
| Scenario (g) | $R \times R, \lambda_A = \lambda_B = 0, MAF_A = MAF_B = 0.4$ | | | |
| 0.05 | 0.05004 | 0.0491 | 0.05035 | 0.05025 |
| 0.005 | 0.00491 | 0.00517 | 0.00507 | 0.00495 |
| 0.0005 | 0.00051 | 0.00056 | 0.00046 | 0.00054 |
| Scenario (h) | $R \times R, \lambda_A = 0.2, \lambda_B = 0, MAF_A = MAF_B = 0.4$ | | | |
| 0.05 | 0.05543 | 0.04863 | 0.18046 | 0.05312 |
| 0.005 | 0.00576 | 0.0051 | 0.04005 | 0.0053 |
| 0.0005 | 0.00071 | 0.00044 | 0.00815 | 0.00062 |

Figure 1 shows power curves for the UMPU test, the PLINK and the LRT tests in the same scenarios as in Table 3. The simulation settings are the same as before, except for the interaction effect which was varied to estimate power curves. As can be seen, the LRT is often the most powerful of all tests, especially in the absence of main effects (scenario d). However, it can be very one–sided (as in scenarios a and c), and can perform quite poorly compared to the UMPU test with less common variants with significant main effects (scenario a). The PLINK and UMPU tests are good at capturing both effect sizes, and the UMPU has more power than PLINK across the board.

**Figure 1.**
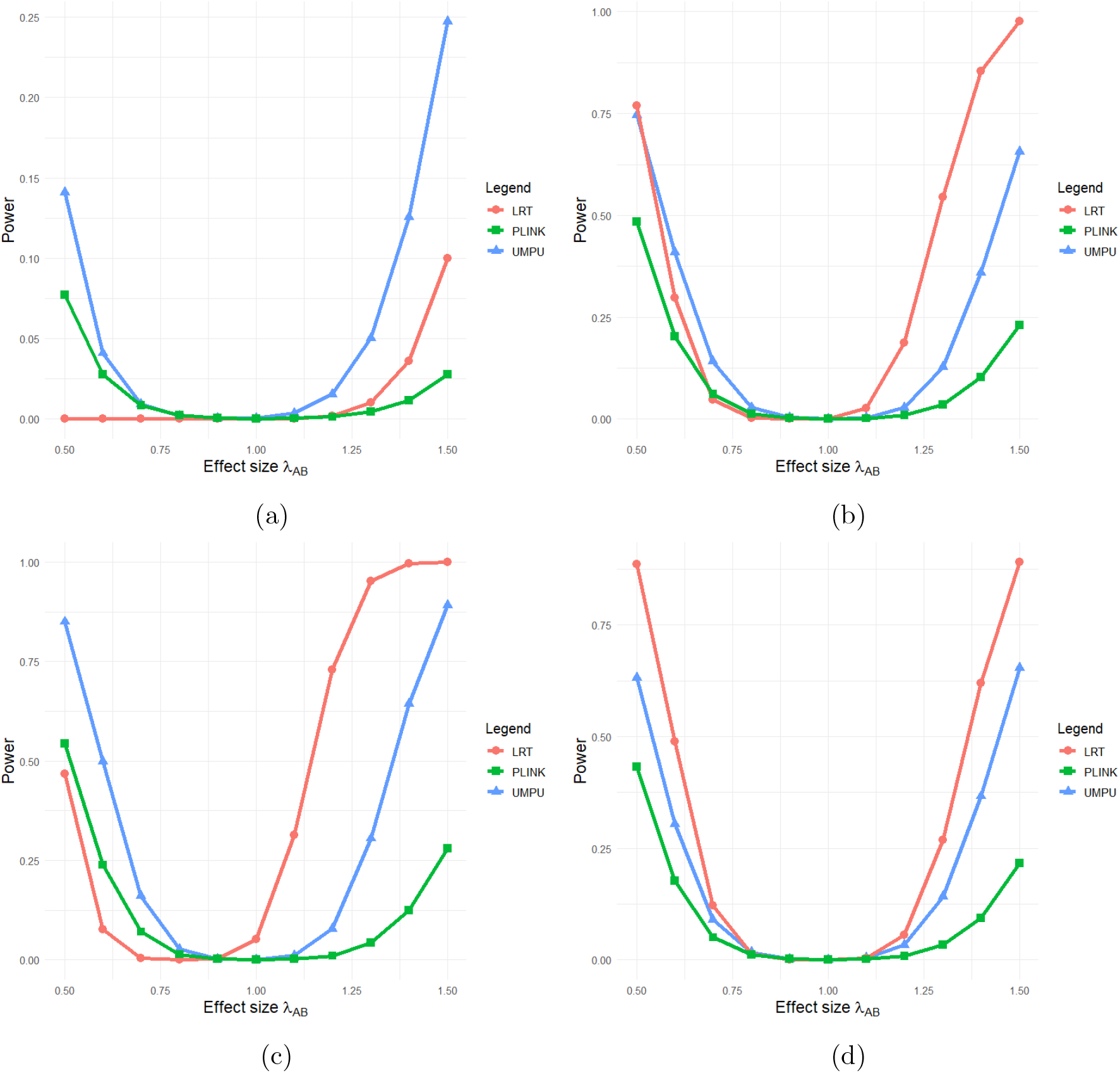
Power curves over interaction effect size, in the following situations: (a) main effect in A; (b) main effects in opposite directions A (0.2) and B (−0.1) (c) effects in the same direction A (0.1) and B (0.2); (d) no main effects. The MAFs for A and B are 0.1 and 0.2, respectively, in all cases except for (a), where the MAF = 0.025 for both. Simulations assume an additive × additive model, and study sizes are as given in Table 3

For scenarios involving dominant and recessive mechanisms of action, the power curves are given in Figure 2. The first thing to notice is that the specific UMPU test is more powerful than the misspecified LRT on at least part of the range of effect sizes. For recessive×recessive with no main effects, the specific UMPU test is significantly more powerful than all the other tests (scenario g). This suggests that when testing recessive mechanisms where the signal is driven by a small number of samples in the study, using a specific test could make a big difference. Between the unbiased tests, the power ranking is consistent throughout all scenarios and ranges, namely PLINK < UMPU < specific UMPU.

**Figure 2.**
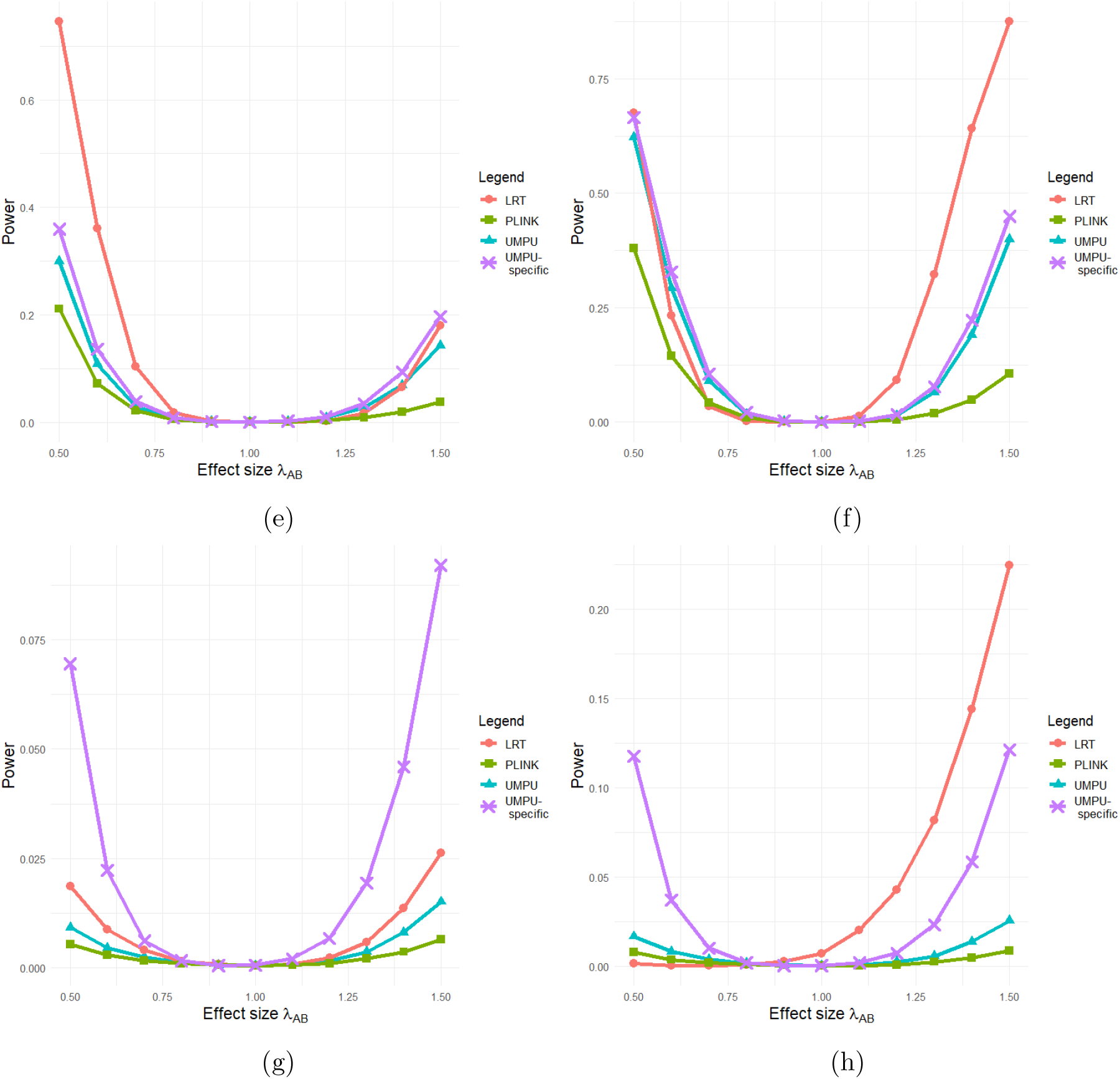
Power curves over interaction effect size, in the following situations: (e) D × D with negative main effect in B only; (f) D × A with main effects in opposite directions; (g) R × R with no main effects; (h) R × R with positive effect in A only. Other settings are as given in Table 4

## 6 Genetic interaction scan for IBD in the UK Biobank

As a real data application, we look for evidence of interaction effects between variants previously reported to be associated with inflammatory bowel disease (IBD). We take the entire list of 232 hits from Liu et al. (2015) and identify them based on their physical locations from the UK Biobank whole genome sequencing data. Part of the motivation for doing this is provided by Evangelou (2018) who concludes that it is rare to find genetic interactions without main effects.

All variants are common (MAF above 0.01) and all pass quality control checks. Cases are obtained by applying the filter for diagnostic codes K50 & K51 (Wu et al., 2023), resulting in 7,051 individuals. Controls are matched to cases on 10 principal components computed from whole genome sequencing, age and sex. We include 28,184 controls corresponding to a 4:1 matching ratio. For each variant, we estimate the MAF based on the control subset rather than for the entire UK Biobank population, to reflect the substructure in our matched cases and controls. To avoid linked variants we compute *R*^2^ for any pair on the same chromosome and filter out pairs with *R*^2^ > 0.0001 (186 in total). Note that this is not the same as LD pruning because we are not dropping entire variants, only pairs that are in LD. We compute the three tests we have been considering – the UMPU, LRT, and PLINK– for each pair of variants in the filtered list, for a total of 25,239 tests involving 226 unique variants.

From Figure 3 we can see that type I errors are well controlled by the UMPU and PLINK tests, and very liberal for the LRT. Given that variants in LD have been excluded, the inflated type I errors for the LRT are likely caused by the presence of main effects. There is generally good agreement between UMPU and PLINK test statistics, with a correlation of 89.5%. Table S1 in the Supplementary Material lists the top pairs, including information on both variants and testing results. None of the variant pairs are individually significant after Bonferroni correction (threshold 2 × 10^−6^), although the top hit comes close. This echoes previous results in Aleknonytė-Resch et al. (2020), who use a case–only design with an otherwise similar search setup. Notably, our smallest p–value is slightly lower than the smallest p–value they found, despite that study using the largest available dataset of IBD (29,524 patients). We then proceed to look for evidence of aggregate effects, by taking the set of pairs that are significant at the more exploratory level of 0.001 and running the list of (unique) gene names that either variant is annotated to through the online gene ontology (Ashburner et al., 2000; The Gene Ontology Consortium et al., 2023). For the subset of genes that were uniquely mapped, a search through the PANTHER database of biological processes (Mi et al., 2019; Thomas et al., 2022) returned two significant ontological terms, the positive regulation of leukocyte activation, and its parent term positive regulation of cell activation. Both pathways were significant after Bonferroni correction of a Fisher’s exact overrepresentation test, with the leukocite activation having a fold enrichment of 13.83 and an adjusted p-value of 0.032. IBD is known to have an inflammatory genetic basis, and leukocyte activity in particular has been targeted for treatment of IBD (see (Arseneau & Cominelli, 2015; Bamias et al., 2013)). This provides some confirmation that interaction signals that we have detected are biologically meaningful. It also suggests that genetic interaction signals might be of modest size for IBD, compared to main effects (some of which having reported p–values below 10^−50^ in the original GWAS study). Coupled with the high number of possible tests in genotype–genotype scans and hence a high price paid for multiple testing, a more feasible approach for future studies might be to find ways to pool evidence from relatively small effects in order to arrive at biologically actionable conclusions.

**Figure 3.**
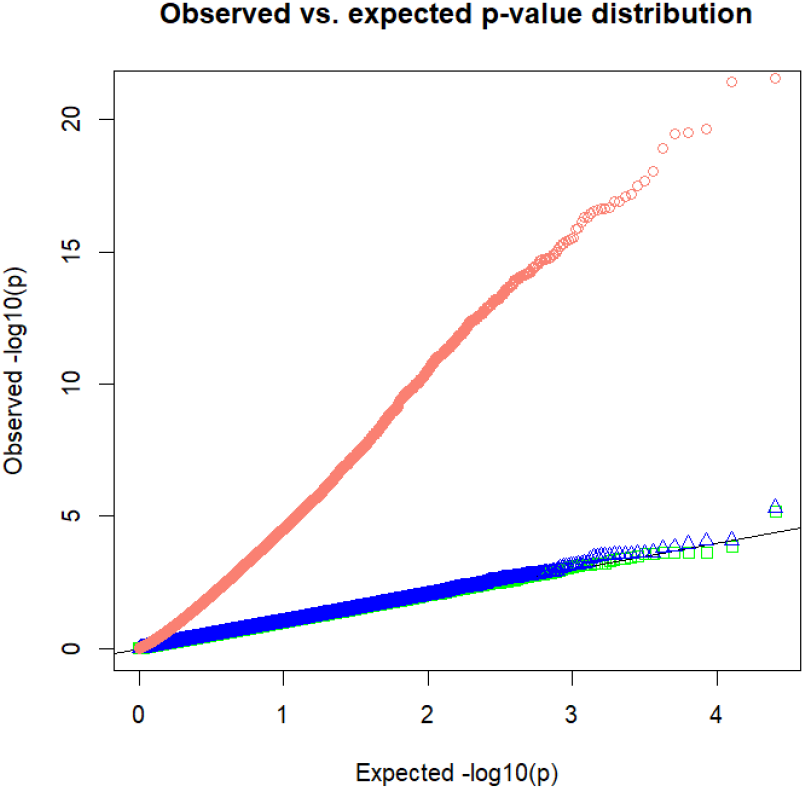
QQ plot of p–values from the targeted scan of interactions in the UK Biobank. Includes LRT (red ‘o’), PLINK (green ‘□’), and UMPU (blue ‘△’).

## 7 Discussion

One of the implications of the UMPU test constructed in this paper is that there is value in control data. Some of the previous tests based on odd ratios either use only cases, or if they do use controls, assume them to be useful only when the two loci are correlated in the population (as discussed by Ueki & Cordell (2012)). However, as we have shown, the sufficient statistics for both main effects and interaction rely on the differenced counts 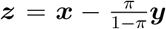, and this assumes no correlation between loci. Hence, only in rare diseases is it acceptable to throw away controls. Otherwise, they play a role in inference, because controls will be impoverished in those genotypes that are enriched in cases, as compared to the general population.

The construction of the conditional UMPU test raises some important points for discussion. Firstly, even though both the single and multi-parameter model tests are most powerful and unbiased, the power profiles for testing the interaction parameter (*γ* or *λ*_*AB*_) can be different. This is because not knowing *λ*_*A*_ and *λ*_*B*_, and thus having to base the test on *U* |***T***, introduces noise into the test statistic. This can cause significant loss of power due to ***T*** containing both information about (*λ*_*A*_, *λ*_*B*_), as well as noise, as we have seen in some of the simulated scenarios. If, by contrast, we had oracle knowledge of the values of nuisance parameters *λ*_*A*_ and *λ*_*B*_, our test statistic would simply be based on the sufficient statistic *U*, and this would be asymptotically equivalent to the single parameter model test (this is clear at least for the case when *λ*_*A*_ and *λ*_*B*_ are both zero). Thus, there is a cost in power to ensure nominal type I error control when testing in the presence of unkown nuisance parameters.

A similar construction could be used to immunize the interaction test statistic against inflated type I error due to the effects of other covariates. However, that will likely incur even greater costs in terms of power and should probably not be used routinely. Matching cases and controls on relevant covariates is probably a better solution to adjust for known confounders. Finally, as a computational note, the test developed in this paper is fast to compute, as it is available in closed form and does not require loops or any other optimization steps.

## 8 Acknowledgments

Support for this research has been provided by the Manitoba Medical Services Foundation. RGR is based at the George & Fay Yee Centre for Healthcare Innovation. Support for CHI is provided by University of Manitoba, Canadian Institutes for Health Research, Province of Manitoba, and Shared Health Manitoba. Bioinformatic support was provided by Travis Haight, from the Statistical Genomics and Bioinformatics Platform (SGB), University of Manitoba. This research has been conducted using the UK Biobank Resource under Application Number 97418.

