## Supplementary Material for "A Most Powerful Test for Gene–Gene Interaction in the Presence of Main Effects"

January 28, 2026

### Proofs

#### Derivation of Constant $k_0$

Using the law of conditional probabilities, we can decompose the prevalence as follows, assuming discrete covariates

$$P(D=1)=\sum_{i,j,x} P(D=1\mid i,j,X)P(i,j,x)=\sum_{i,j,x} f(\boldsymbol{\lambda},\boldsymbol{\beta};i,j,x)P(i,j,x),$$

where the summation is over all covariates and genotypes. If we further assume that covariates are independent of the genotypes at loci $A$ and $B$, we can further simplify

$$P(D=1)=\sum_{i,j,x} f(\boldsymbol{\lambda},\boldsymbol{\beta};i,j,x)\mu_{ij}P(X=x)=\sum_{i,j,x}\pi k_0 e^{x\boldsymbol{\beta}}(1+\lambda_A)^i(1+\lambda_B)^j(1+\lambda_{AB})^{ij}\mu_{ij}P(X=x)$$

$$\Leftrightarrow \pi=\pi k_0\left(\sum_{i,j}(1+\lambda_A)^i(1+\lambda_B)^j(1+\lambda_{AB})^{ij}\mu_{ij}\right)\left(\sum_x e^{x\boldsymbol{\beta}}P(X=x)\right)$$

$$\Rightarrow k_0=\left(\sum_{i,j}(1+\lambda_A)^i(1+\lambda_B)^j(1+\lambda_{AB})^{ij}\mu_{ij}\right)^{-1}\left(\sum_x e^{x\boldsymbol{\beta}}P(X=x)\right)^{-1}.$$

For penetrance model (6) in the main text, this implies

$$k_0(\gamma)=(\mu_{00}+\mu_{01}+\mu_{02}+\mu_{10}+\mu_{20}+\gamma\mu_{11}+\gamma^2(\mu_{12}+\mu_{21})+\gamma^4\mu_{22})^{-1}. \tag{1}$$

For genetic model (7) we can write, using the approximation formulas for small effect sizes,

$$\begin{aligned}\pi=\pi k_0[&\mu_{00}+(1+\lambda_A)\mu_{10}+(1+2\lambda_A)\mu_{20}+(1+\lambda_B)\mu_{01}+(1+2\lambda_B)\mu_{02}\\&+(1+\lambda_A+\lambda_B+\lambda_{AB})\mu_{11}+(1+\lambda_A+2\lambda_B+2\lambda_{AB})\mu_{12}+(1+2\lambda_A+\lambda_B+2\lambda_{AB})\mu_{21}\\&+(1+2\lambda_A+2\lambda_B+4\lambda_{AB})\mu_{22}]\end{aligned}$$

Rearranging and solving for $k_0$ yields

$$k_0=\frac{1}{1+\lambda_A\boldsymbol{a}\cdot\boldsymbol{\mu}+\lambda_B\boldsymbol{b}\cdot\boldsymbol{\mu}+\lambda_{AB}(\boldsymbol{a}\odot\boldsymbol{b})\cdot\boldsymbol{\mu}}. \tag{2}$$

### Derivation of the Likelihood Ratio Test (LRT)

First, we must find the maximum likelihood estimate $\hat{\gamma}$ which maximizes the log-likelihood function (4) in the text. Thus, $\hat{\gamma}$ maximizes

$$\begin{aligned} l(\gamma) &= \varphi_1(\boldsymbol{x},\boldsymbol{y}) + \sum_{i,j\in\{0,1,2\}} x_{ij}\log(f(\gamma;i,j)) + y_{ij}\log(1-f(\gamma;i,j)) \\ &= \varphi_1(\boldsymbol{x},\boldsymbol{y}) + \sum_{i,j\in\{0,1,2\}} x_{ij}[ij\cdot\log(\gamma)+\log(\pi)+\log(k_0(\gamma))] + \sum_{i,j\in\{0,1,2\}} y_{ij}\log(1-\pi k_0(\gamma)\gamma^{ij}), \end{aligned} \tag{3}$$

where $k_0$ is given in 1. To build the LRT for testing $\gamma = 1$ versus a two–sided alternative $\gamma \neq 1$, use the usual test statistic

$$T_{LR} = -2[l(1) - l(\hat{\gamma})] = \sum_{i,j\in\{0,1,2\}} \left[ x_{ij}\log\left(\frac{f(\hat{\gamma};i,j)}{\pi}\right) + y_{ij}\log\left(\frac{1-f(\hat{\gamma};i,j)}{1-\pi}\right)\right]$$

It is easy to show that, for the reduced model ($\gamma = 1$), $k_0(1) = 1$, and

$$\begin{aligned} l(1) &= \varphi_1(\boldsymbol{x},\boldsymbol{y}) + \log(\pi k_0(1))\left(\sum_{i,j\in\{0,1,2\}} x_{ij}\right) + \log(1-\pi k_0(1))\left(\sum_{i,j\in\{0,1,2\}} y_{ij}\right) \\ &= \varphi_1(\boldsymbol{x},\boldsymbol{y}) + n_1\log(\pi) + n_0\log(1-\pi) \end{aligned}$$

Then,

$$\begin{aligned} T_{LR} &= 2\{\varphi_1(\boldsymbol{x},\boldsymbol{y}) + \sum_{i,j\in\{0,1,2\}} x_{ij}[ij\cdot\log(\hat{\gamma})+\log(\pi)+\log(k_0(\hat{\gamma}))] + \sum_{i,j\in\{0,1,2\}} y_{ij}\log(1-\pi k_0(\hat{\gamma})\hat{\gamma}^{ij})\} \\ &\quad - 2\{\varphi_1(\boldsymbol{x},\boldsymbol{y}) + n_1\log(\pi) + n_0\log(1-\pi)\} \\ &= 2\sum_{i,j\in\{0,1,2\}} \{x_{ij}ij\log\hat{\gamma} + y_{ij}\log\left[1-\pi\hat{\gamma}^{ij}k_0(\hat{\gamma})\right]\} + 2n_1\log k_0(\hat{\gamma}) - 2n_0\log(1-\pi) \end{aligned}$$

This LRT can be argued to be UMPU, and a sketch of the proof is as follows. By approximating all terms in 3 involving $\gamma$ at the first order around 1 (or $\delta = 0$, where $\gamma = 1+\delta$) we can write the log pmf $\log P(\boldsymbol{x},\boldsymbol{y};\delta) \approx h(\boldsymbol{x},\boldsymbol{y}) + g(\delta) + \delta T(\boldsymbol{x},\boldsymbol{y})$, where the sufficient statistic $T(\boldsymbol{x},\boldsymbol{y})$ is some linear combination of $x_{ij}$ and $y_{ij}$. Thus, $T$ is approximately normal for large sample sizes.Thus, our distribution $\boldsymbol{x},\boldsymbol{y}$ is in the single parameter exponential family dependent on $\delta$ (recall that other parameters, such as $\mu_{ij}$ are treated as fixed constants). According to Bhattacharya & Burman (2016) (p. 137) since $T$ has a symmetrical distribution, a symmetric level $-\ \alpha$ test is UMPU. Note that a test based on $T(\boldsymbol{x},\boldsymbol{y})$ is technically a score test. However, it is known that LRT and the score test are asymptotically equivalent Engle (1984); Gałecki & Burzykowski (2013). Thus, we can conclude that the LRT is UMPU for the two-sided testing of $\gamma = 1$ .

### Approximating the expression $\log(1-\pi(1+\delta))$

Here, $\pi$ is the prevalence of a condition, and $\delta$ is a value close to zero. We have

$$\begin{aligned}\log(1-\pi(1+\delta)) = \log(1-\pi-\pi\delta) &= \log(\frac{1-\pi-\pi\delta}{1-\pi}) + \log(1-\pi)\\ &= \log(1-\frac{\pi}{1-\pi}\delta) + \log(1-\pi) \approx -\frac{\pi}{1-\pi}\delta + \log(1-\pi).\end{aligned}$$

### Approximate distributions of vectors x and y

Using expression (9)for $p_{ij}$, we can write

$$\begin{aligned}\text{Var}[X_{ij}] &= n_1 p_{ij}(1-p_{ij}) = n_1\mu_{ij}\left(1+\lambda_A(i-\boldsymbol{a}^\intercal\boldsymbol{\mu})+\lambda_B(j-\boldsymbol{b}^\intercal\boldsymbol{\mu})\right)\left[1-\mu_{ij}\left(1+\lambda_A(i-\boldsymbol{a}^\intercal\boldsymbol{\mu})+\lambda_B(j-\boldsymbol{b}^\intercal\boldsymbol{\mu}\right.\right.\\ &\approx n_1\mu_{ij}\{(1+\lambda_A(i-\boldsymbol{a}^\intercal\boldsymbol{\mu})+\lambda_B(j-\boldsymbol{b}^\intercal\boldsymbol{\mu}))-\mu_{ij}\left(1+2\lambda_A(i-\boldsymbol{a}^\intercal\boldsymbol{\mu})+2\lambda_B(j-\boldsymbol{b}^\intercal\boldsymbol{\mu})\right)\}\\ &= n_1\mu_{ij}(1-\mu_{ij})+\lambda_A n_1\mu_{ij}(1-2\mu_{ij})(i-\boldsymbol{a}^\intercal\boldsymbol{\mu})+\lambda_B n_1\mu_{ij}(1-2\mu_{ij})(j-\boldsymbol{b}^\intercal\boldsymbol{\mu});\quad \text{and}\end{aligned}$$

$$\begin{aligned}\text{Cov}[X_{i_1j_1}, X_{i_2j_2}] = -\, n_1 p_{i_1j_1}p_{i_2j_2} &\approx -n_1\mu_{i_1j_1}\mu_{i_2j_2}[1+\lambda_A(i_1+i_2)+\lambda_B(j_1+j_2)]\\ &+ 2n_1\mu_{i_1j_1}\mu_{i_2j_2}\lambda_A\boldsymbol{a}^\intercal\boldsymbol{\mu} + 2n_1\mu_{i_1j_1}\mu_{i_2j_2}\lambda_B\boldsymbol{b}^\intercal\boldsymbol{\mu}.\end{aligned}$$

To consolidate these expressions in matrix form, it can be verified that the off-diagonal part of the covariance matrix of $\boldsymbol{x}$, $\Sigma_x$ is the same as the off-diagonal part of the matrix $-n_1\boldsymbol{\mu}\boldsymbol{\mu}^\intercal - n_1\lambda_A(A+A^\intercal-2\boldsymbol{a}^\intercal\boldsymbol{\mu}J)\odot(\boldsymbol{\mu}\boldsymbol{\mu}^\intercal) - n_1\lambda_B(B+B^\intercal-2\boldsymbol{b}^\intercal\boldsymbol{\mu}J)\odot(\boldsymbol{\mu}\boldsymbol{\mu}^\intercal)$, where square matrices $A$ and $B$ of size $9\times 9$ are defined as $A = (\boldsymbol{a}\quad\boldsymbol{a}\quad\ldots\quad\boldsymbol{a})$, $B = (\boldsymbol{b}\quad\boldsymbol{b}\quad\ldots\quad\boldsymbol{b})$ and $J = \mathbf{1}\mathbf{1}^\intercal$ is the matrix of ones. Note that the product $\odot$ similarly applies to matrices. The diagonal of $\Sigma_x$ has a few extra terms, but we can write everything as one expression:

$$\begin{aligned}\Sigma_x = n_1 D_{\boldsymbol{\mu}} - n_1\boldsymbol{\mu}\boldsymbol{\mu}^\intercal &+ n_1\lambda_A[D_{\boldsymbol{\mu}\odot(\boldsymbol{a}-\boldsymbol{a}^\intercal\boldsymbol{\mu})} - (A+A^\intercal-2\boldsymbol{a}^\intercal\boldsymbol{\mu}J)\odot(\boldsymbol{\mu}\boldsymbol{\mu}^\intercal)]\\ &+ n_1\lambda_B[D_{\boldsymbol{\mu}\odot(\boldsymbol{b}-\boldsymbol{b}^\intercal\boldsymbol{\mu})} - (B+B^\intercal-2\boldsymbol{b}^\intercal\boldsymbol{\mu}J)\odot(\boldsymbol{\mu}\boldsymbol{\mu}^\intercal)],\end{aligned} \tag{4}$$

where the notation $D_{\boldsymbol{v}}$ denotes the diagonal matrix with diagonal $\boldsymbol{v}$ and zero everywhere else. Other (potentially) useful facts about the matrices used in 4: $A$ can be written $A = D_{\boldsymbol{a}}J$; as well, $\boldsymbol{\mu}\boldsymbol{\mu}^\intercal = D_{\boldsymbol{\mu}}JD_{\boldsymbol{\mu}}$. Furthermore, for any square matrix $M$, we have $M\odot(\boldsymbol{\mu}\boldsymbol{\mu}^\intercal) = D_{\boldsymbol{\mu}}MD_{\boldsymbol{\mu}}$. For vector $\boldsymbol{y}$ we have

$$\begin{aligned}\text{Var}[Y_{ij}] &= n_0 q_{ij}(1-q_{ij})\\ &\approx n_0\mu_{ij}[1-\frac{\pi}{1-\pi}\left(\lambda_A(i-\boldsymbol{a}^\intercal\boldsymbol{\mu})+\lambda_B(j-\boldsymbol{b}^\intercal\boldsymbol{\mu})\right)][1-\mu_{ij}+\mu_{ij}\frac{\pi}{1-\pi}\left(\lambda_A(i-\boldsymbol{a}^\intercal\boldsymbol{\mu})+\lambda_B(j-\boldsymbol{b}^\intercal\boldsymbol{\mu})\right)]\\ &= n_0\mu_{ij}(1-\mu_{ij})[1-\frac{\pi}{1-\pi}\left(\lambda_A(i-\boldsymbol{a}^\intercal\boldsymbol{\mu})+\lambda_B(j-\boldsymbol{b}^\intercal\boldsymbol{\mu})\right)][1+\frac{\mu_{ij}}{1-\mu_{ij}}\frac{\pi}{1-\pi}\left(\lambda_A(i-\boldsymbol{a}^\intercal\boldsymbol{\mu})+\lambda_B(j-\right.\\ &\approx n_0\mu_{ij}(1-\mu_{ij})\{1+\frac{\pi}{1-\pi}[\lambda_A(i-\boldsymbol{a}^\intercal\boldsymbol{\mu})+\lambda_B(j-\boldsymbol{b}^\intercal\boldsymbol{\mu})]\left(\frac{\mu_{ij}}{1-\mu_{ij}}-1\right)\}\\ &= n_0\mu_{ij}(1-\mu_{ij})+n_0\mu_{ij}(2\mu_{ij}-1)\frac{\pi}{1-\pi}[\lambda_A(i-\boldsymbol{a}^\intercal\boldsymbol{\mu})+\lambda_B(j-\boldsymbol{b}^\intercal\boldsymbol{\mu})]\quad\text{and}\end{aligned}$$

$$\begin{aligned}\text{Cov}[Y_{i_1j_1}, Y_{i_2j_2}] &= -n_0 q_{i_1j_1}q_{i_2j_2}\\ &\approx -n_0\mu_{i_1j_1}\mu_{i_2j_2}\left[1-\lambda_A\frac{\pi}{1-\pi}(i_1+i_2-2\boldsymbol{a}^\intercal\boldsymbol{\mu})-\lambda_B\frac{\pi}{1-\pi}(j_1+j_2-2\boldsymbol{b}^\intercal\boldsymbol{\mu})\right].\end{aligned}$$

We can similarly write the covariance of $\boldsymbol{y}$ in matrix form as

$$\begin{aligned}\Sigma_y = n_0 D_{\boldsymbol{\mu}} - n_0 \boldsymbol{\mu}\boldsymbol{\mu}^\intercal - n_0 \lambda_A \frac{\pi}{1-\pi}[D_{\boldsymbol{\mu}\odot(\boldsymbol{a}-\boldsymbol{a}^\intercal\boldsymbol{\mu})} - (A + A^\intercal - 2\boldsymbol{a}^\intercal\boldsymbol{\mu}J) \odot (\boldsymbol{\mu}\boldsymbol{\mu}^\intercal)] \\ - n_0 \lambda_B \frac{\pi}{1-\pi}[D_{\boldsymbol{\mu}\odot(\boldsymbol{b}-\boldsymbol{b}^\intercal\boldsymbol{\mu})} - (B + B^\intercal - 2\boldsymbol{b}^\intercal\boldsymbol{\mu}J) \odot (\boldsymbol{\mu}\boldsymbol{\mu}^\intercal)],\end{aligned} \tag{5}$$
